# Genomic analysis of the diversity, antimicrobial resistance and virulence potential of clinical *Campylobacter jejuni* and *Campylobacter coli* strains from Chile

**DOI:** 10.1101/2020.07.03.182048

**Authors:** Veronica Bravo, Assaf Katz, Lorena Porte, Thomas Weitzel, Carmen Varela, Narjol Gonzalez-Escalona, Carlos J. Blondel

## Abstract

*Campylobacter jejuni* and *Campylobacter coli* are the leading cause of human gastroenteritis in the industrialized world and an emerging threat in developing countries. The incidence of campylobacteriosis in South America is greatly underestimated, mostly due to the lack of adequate diagnostic methods. Accordingly, there is limited genomic and epidemiological data from this region. In the present study, we performed a genome-wide analysis of the genetic diversity, virulence, and antimicrobial resistance of the largest collection of clinical *C. jejuni* and *C. coli* strains from Chile available to date (n=81), collected in 2017-2019 in Santiago, Chile. This culture collection accounts for over a third of the genome sequences available of clinical strains from South America. cgMLST analysis identified high genetic diversity as well as 13 novel STs and alleles in both *C. jejuni* and *C. coli*. Pangenome and virulome analyses showed a differential distribution of virulence factors, including both plasmid and chromosomally encoded T6SSs and T4SSs. Resistome analysis predicted widespread resistance to fluoroquinolones, but low rates of erythromycin resistance. This study provides valuable genomic and epidemiological data and highlights the need for further genomic epidemiology studies in Chile and other South American countries to better understand molecular epidemiology and antimicrobial resistance of this emerging intestinal pathogen.

**AUTHOR SUMMARY:** *Campylobacter* is the leading cause of bacterial gastroenteritis worldwide and an emerging and neglected pathogen in South America. In this study, we performed an in-depth analysis of the genome sequences of 69 *C. jejuni* and 12 *C. coli* clinical strains isolated from Chile, which account for over a third of the sequences from clinical strains available from South America. We identified a high genetic diversity among *C. jejuni* strains and the unexpected identification of clade 3 *C. coli* strains, which are infrequently isolated from humans in other regions of the world. Most strains harbored the virulence factors described for *Campylobacter*. While ∼40% of strains harbored mutation in the *gyrA* gene described to confer fluoroquinolone resistance, very few strains encoded the determinants linked to macrolide resistance, currently used for the treatment of campylobacteriosis. Our study contributes to our knowledge of this important foodborne pathogen providing valuable data from South America.

## INTRODUCTION

Campylobacteriosis caused by *Campylobacter jejuni* and *Campylobacter coli* has emerged as an important public health problem and is the most common bacterial cause of human gastroenteritis worldwide [1,2]. It is usually associated with a self-limiting illness characterized by diarrhea, nausea, abdominal cramps and bloody stools [1]. Nevertheless, extraintestinal manifestations might cause long-term complications such as Miller-Fisher or Guillain-Barré syndrome [1]. Despite the self-limiting course, certain populations such as children under 5 years of age, elderly patients, and immunocompromised patients might suffer severe infections and require antimicrobial treatment [3]. In this context, the World Health Organization (WHO) has included *Campylobacter* as a high priority pathogen due to the worldwide emergence of strains with high level fluoroquinolone resistance [4].

In South America, there is limited data available on the prevalence of *C. jejuni* and *C. coli* in comparison to other enteric bacterial pathogens [1,5,6]. In Chile, the National Laboratory Surveillance Program of the Public Health Institute has reported an incidence rate of campylobacteriosis of 0.1 to 0.6 cases per 100,000 persons [7]. This is low compared to high-income countries, where incidence rates can reach two orders of magnitude higher [1]. Nevertheless, recent reports suggest that the burden of campylobacteriosis in Chile has been greatly underestimated due to suboptimal diagnostic protocols [7]. Indeed, a recent report suggests that *Campylobacter* spp. is emerging as the second most prevalent bacterial cause of human gastroenteritis in Chile [8].

Only few reports have analyzed the genetic diversity and antimicrobial resistance profiles of clinical human strains of *Campylobacter* in Chile [9–13]. Furthermore, these studies have either used a limited number of strains [13] or lacked the resolution provided by whole genome sequence-based typing methods [9–12]. Therefore, larger genomic epidemiology studies are needed in Chile and more broadly in South America.

We recently made available the whole genome sequences of 69 *C. jejuni* and 12 *C. coli* human clinical strains isolated from patients with an enteric infection visiting Clínica Alemana in Santiago, Chile, during a 2-year period (2017-2019) [14,15]. This represents the largest genome collection of clinical *Campylobacter* strains from Chile and one of the few collections available from South America. In the present study, we performed a genome-wide analysis of the genetic diversity, virulence potential, and antimicrobial resistance profiles of these strains. Importantly, we identified a high genetic diversity of both *C. jejuni* and *C. coli* strains, including 13 novel sequence types (STs) and alleles. Resistome analysis predicted widespread resistance to fluoroquinolones, but decreased resistance to tetracycline in comparison to the reported resistance from other regions of the world. In addition, we also describe differential distribution of virulence factors among strains, including plasmids and chromosomally encoded Type 6 and Type 4 Secretion Systems (T6SS and T4SS respectively). This work provides valuable epidemiological and genomic data of the diversity, virulence and resistance profiles of this neglected foodborne pathogen.

## METHODS

### Genome data set, sequencing and reannotation

The genome sequence dataset analyzed in this study is listed in **Table S1** and includes our recently published collection of human clinical strains of *C. jejuni* and *C. coli* from Chile [14,15]. We additionally sequenced *C. jejuni* strain CFSAN096303. This strain was sequenced as described in [14] using the NextSeq sequencer (Illumina) obtaining an estimated genome average coverage of 90X. The genome was assembled using the CLC Genomics Workbench v9.5.2 (Qiagen) following the previously used pipeline [14] and was deposited under accession number WXZQ01000000. FASTA files of all draft genomes were reordered to the reference genomes of the *C. coli* aerotolerant strain OR12 (accID GCF_002024185.1) and *C. jejuni* strain NCTC 11168 (accID GCF_000009085.1) using the Mauve Contig Mover (MCM) from the Mauve package [16]. The ordered genomes were annotated using Prokka [17] with the same reference files mentioned above and by forcing GenBank compliance. Draft genome sequence reannotations and sequence FASTA files are available as a supplementary dataset on Zenodo https://doi.org/10.5281/zenodo.3925206.

### Multi-Locus Sequence Typing (MLST) and core genome MLST (cgMLST) analysis

Genome assemblies were mapped against an MLST scheme based on seven housekeeping genes [18] and the 1,343-locus cgMLST scheme [19], using RIDOM SeqSphere software (Münster, Germany) [20]. The following GenBank (GCA) or RefSeq (GCF) genome assemblies were used as reference genomes: *Campylobacter hepaticus HV10* (GCF_001687475.2), *Campylobacter upsaliensis DSM 5365* (GCF_000620965.1), *Campylobacter lari RM2100* (GCF_000019205.1), *Campylobacter jejuni NCTC 11168* (GCF_000009085.1*), C. jejuni doylei 269*.*97* (GCA_000017485.1), aerotolerant *Campylobacter coli OR12* (GCF_002024185.1), *Campylobacter coli BIGS0008* (GCA_000314165.1), *Campylobacter coli 76339* (GCA_000470055.1), and *Campylobacter coli H055260513* (GCA_001491555.1) were used as reference genomes. Phylogenetic comparison of the 81 *Campylobacter* genome sequences was performed using a neighbor-joining tree based on a distance matrix of the core genomes of all strains. Sequence Types (STs) and Clonal Complexes (CCs) were determined after automated allele submission to the *Campylobacter* PubMLST server [18]. Comparative cgMLST analysis with *C. jejuni* and *C. coli* clinical strains worldwide was performed with genomes available at the NCBI database. Annotations and visualizations were performed using iTOL v.4 [21].

### Virulome, resistome and comparative genomic analysis

A local BLAST database was constructed using FASTA files of all sequenced genomes and the makeblastdb program from BLAST [22], which was screened for the nucleotide sequence of known pathogenicity genes using BLASTn (version 2.8.1) [22]. A 90% sequence length and 90% identity threshold were used to select positive matches. The FASTA file containing the nucleotide sequence of virulence genes screened can be found at https://doi.org/10.5281/zenodo.3925206. The C257T mutation in *gyrA* and mutations A2058C and A2059G in the genes for 23S RNA were searched manually by BLAST and sequences were aligned to the reference gene from strain *Campylobacter jejuni* ATCC 33560 using ClustalX [23]. Antimicrobial resistance genes were screened in each *Campylobacter* genome using the ABRicate pipeline [24], using the Resfinder [25], CARD [26], ARG-ANNOT [27] and NCBI ARRGD [28] databases. BLAST-based genome comparisons and visualization of genetic clusters were performed using EasyFig [29].

### Pangenome and plasmid screening analysis

Annotated genome sequences were compared using Roary [30], setting a minimum percentage identity of 90% for BLASTp. Pangenome visualization was performed using roary_plots.py. Additionally, the file gene_presence_absence.csv produced by Roary was analyzed using a spreadsheet program to find genes that are shared by subgroups of isolates. Venn diagrams were constructed using the tool available at http://bioinformatics.psb.ugent.be/webtools/Venn/. Plasmids screening was performed on FASTQ files using PlasmidSeeker [31], based on the default bacterial plasmid database. The inhouse database of known plasmids from *C. coli* or *C. jejuni* and the output table for the presence/abscence of gene content in the Roary analysis can be found in https://doi.org/10.5281/zenodo.3925206. Pangenome BLAST analysis was performed using GView (https://server.gview.ca/) [32].

## RESULTS

## Phylogenetic analysis of clinical Campylobacter strains

To gain insight into the diversity of Chilean clinical *C jejuni* and *C. coli* jejuni strains, we performed a cgMLST analysis using the typing scheme recently developed by Cody *et al* [19]. cgMLST analysis showed high genetic diversity among the 69 *C. jejuni* strains, as shown by the 14 CCs and 31 distinct STs identified (**Fig 1**). Almost 93% (n=64) of the *C. jejuni* strains belonged to a previously described CC for this specie. Among our strains, CC-21 was the most common (37.6%) and diverse (5 distinct STs) (**Fig 1, S1 Table**). ST-1359 was the most prevalent ST of CC-21 (48%) and of all *C. jejuni* strains in this study (17.4%) (**Fig 1 and S1 Table**). In addition to CC-21, we identified high rates of CC-48 (13%) and CC-52 (7.3%). ST-475 from CC-48 was the second most prevalent ST (11.6%) identified (**Fig 1**).

**Fig 1.**
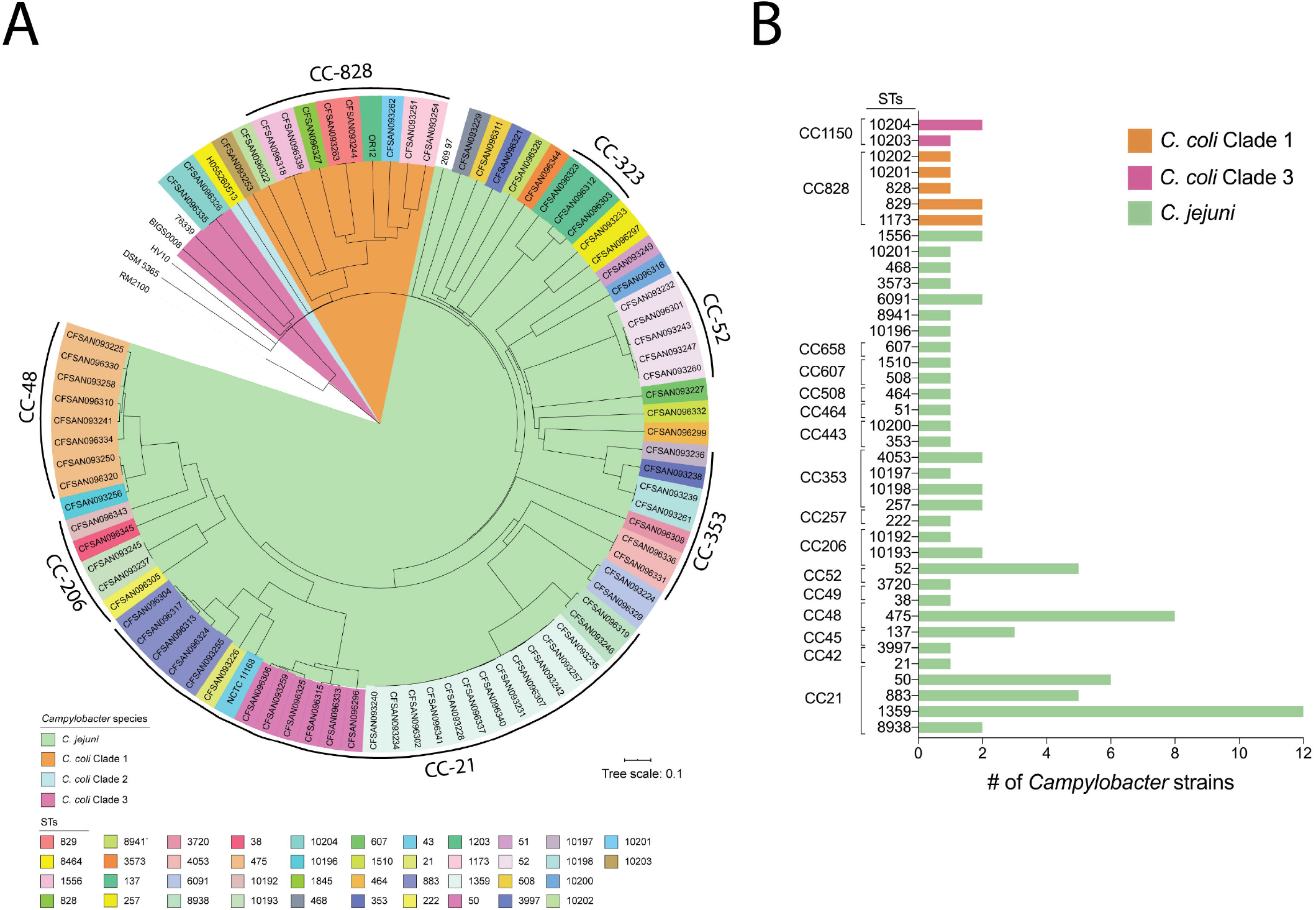
Phylogenetic analysis of clinical *C. jejuni* and *C. coli* strains from Chile. Phylogenetic tree is based on cgMLST performed with RIDOM SeqSphere^+^ and visualized by iTOL. STs are shown in colored boxes for each strain, and tree branches are color coded to highlight *C. jejuni* and *C. coli* strains from clades 1, 2 and 3. ST and CC frequency distribution of clinical *C. jejuni* and *C. coli* strains from this study are detailed in the right panel.

cgMLST analysis of the 12 *C. coli* strains from our collection revealed that 83% (10/12) of the strains belonged to clade 1, while 17% (2/12) belonged to clade 3 and none to clade 2. Most clade 1 strains (8/10) belonged to CC-828 and included four previously described STs (828, 829, 1173, and 1556) and one novel ST (10201). Only one strain from clade 1 belonged to the usually common CC-1150 (strain CFSAN093253 from ST-10203), and another strain could not be assigned to any clonal complex (strain CFSAN096322 from ST-10202). Both clade 3 strains belonged to ST-10204.

A global cgMLST analysis comparing our 81 *Campylobacter* genomes with other 1631 *C. jejuni* and 872 *C. coli* genomes available from the NCBI database was performed. The strains of our collections from Central Chile showed great diversity and did not group with any existing clusters (**Fig 2A)**. The reconstructed phylogenetic trees and associated metadata of clinical strains from Chile in the global context of *C. jejuni* and *C. coli* sequenced strains can be interactively visualized on Microreact at https://microreact.org/project/VbEQsZtQD. Analysis of the genotype frequencies of clinical *Campylobacter* strains from Chile in comparison to other countries, showed a distribution of CCs more similar to countries from North America (United States) and Europe (United Kingdom) than from other countries from the region such as Peru, Brazil and Uruguay (**Fig 2B**).

**Fig 2.**
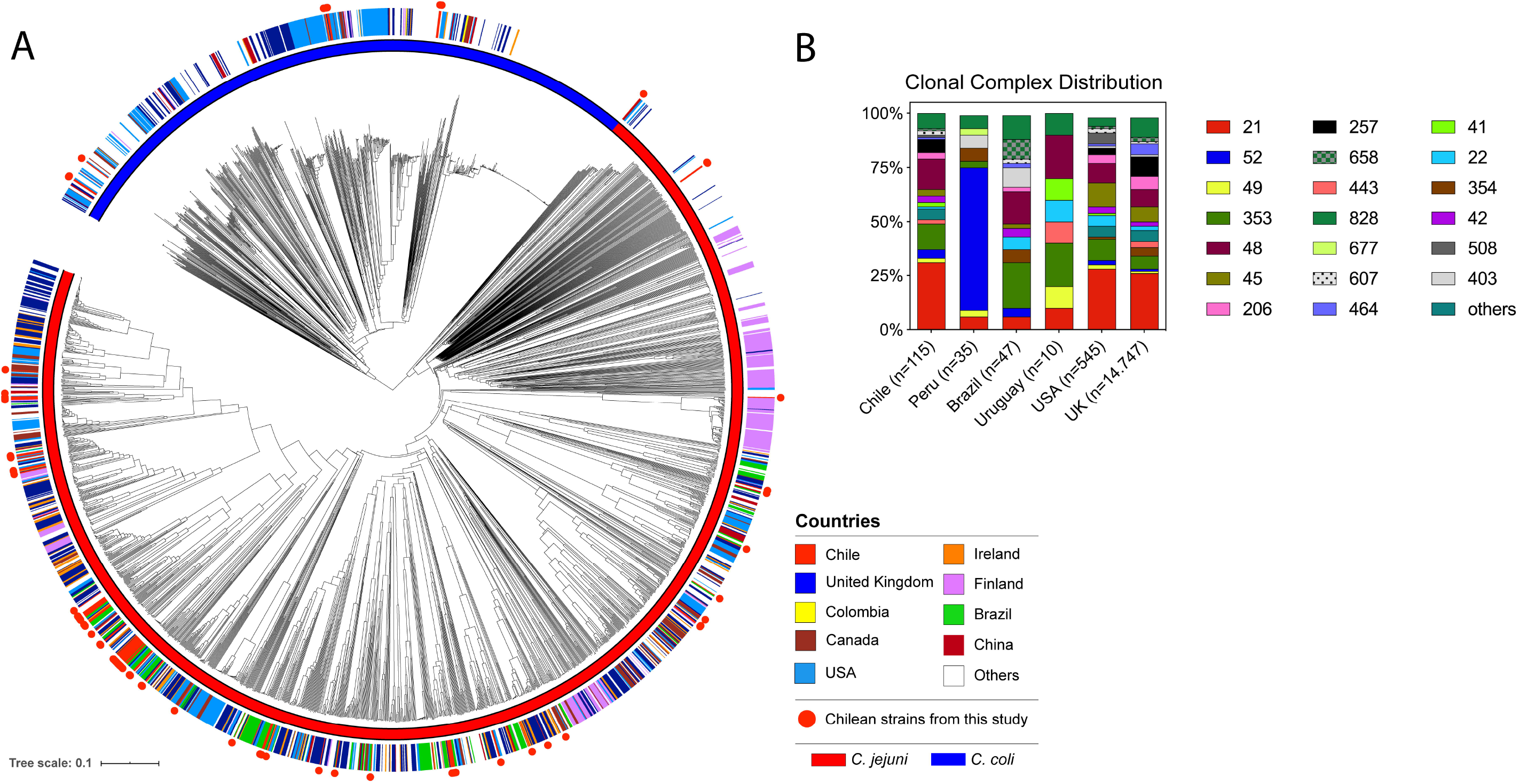
Global cgMLST analysis of clinical *C. jejuni* and *C. coli* strains. (A) Phylogenetic tree is based on cgMLST performed with RIDOM SeqSphere^+^ and visualized by iTOL. Countries of isolation are shown as colored strip and strains from this study are highlighted in red circles. A blue or red color strip highlights the species of each isolate. (B) Clonal complex frequency of clinical *Campylobacter* strains deposited in the pubMLST database separated per country of isolation.

### Pangenome analysis of Campylobacter strains

To identify the genetic elements that are shared and distinct among the Chilean *C. jejuni* and *C. coli* strains, we performed pangenome analysis of the 81 strains from both species (**Fig 3**). The analysis identified a pangenome of 6609 genes for both species and identified the core genes shared between *C. jejuni* and *C. coli* (531 genes) and the core genome of each species (1300 genes for *C. jejuni* and 1207 genes for *C. coli*). Our analysis showed an open pangenome as the number of genes increases with the addition of new genomes. The core genome decreased reaching a plateau at approximately 20 genomes (**Fig 2C**). The number of new and unique genes identified also reached a plateau after 20 genomes (**Fig 2D**).

**Fig 3.**
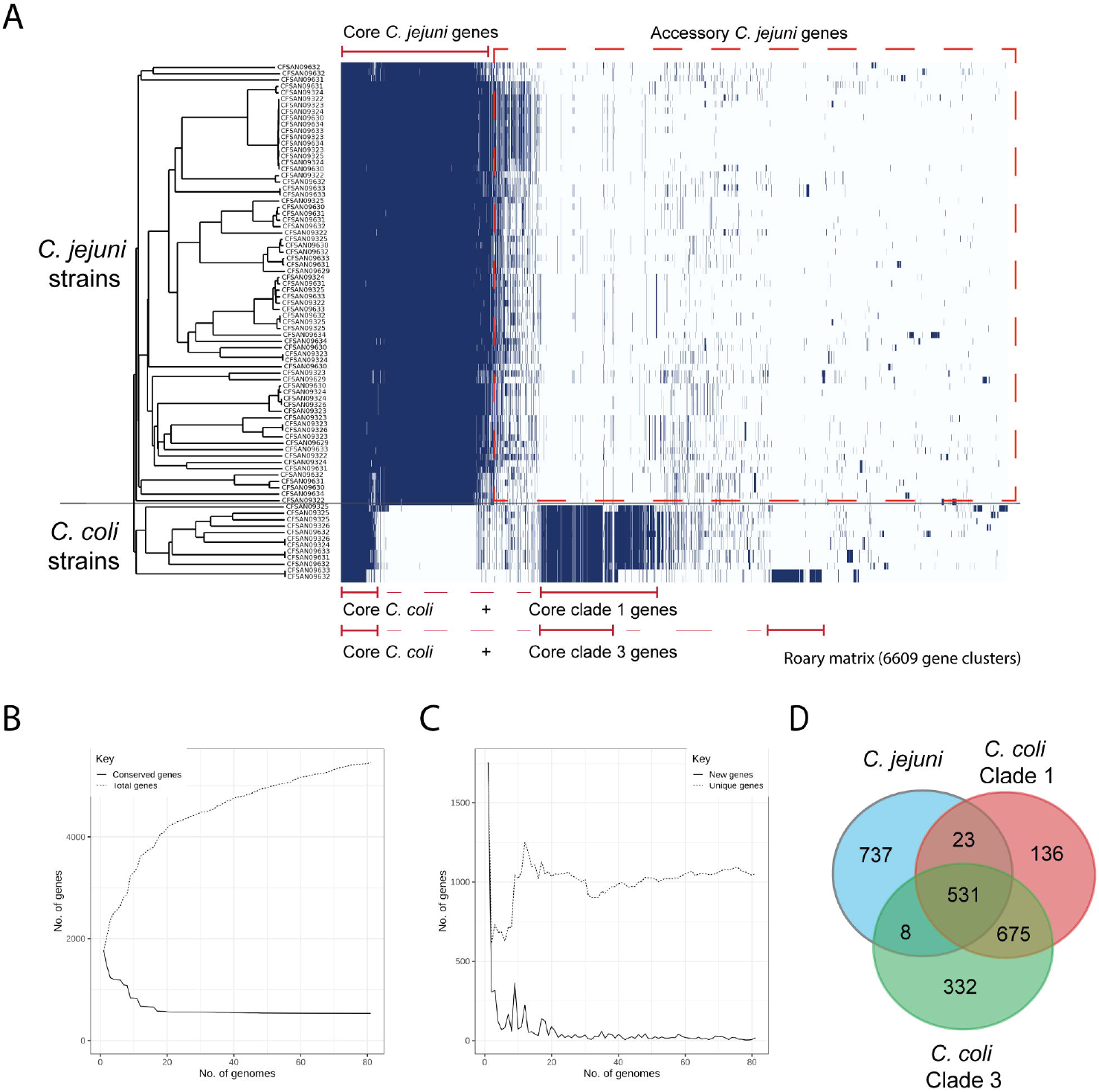
Visualization of the pangenome of the *C. jejuni* and *C. coli* strains. Pangenome analysis was performed using Roary. Core and accessory genomes are highlighted with red and black brackets, respectively. cgMLST phylogenetic tree of Fig 1A is shown on the left. B) The graph shows the number of genes in the core and pan genomes (continuous and discontinuous lines respectively) as increasing number of genomes are considered in random order. C) The number of genes unique to a genome (discontinuous line) and new genes not found in previously analyzed strains (continuous line) as increasing number of genomes are considered in random order. D) A Venn diagram highlighting the number of core genes present in *C. jejuni* and *C. coli* strains from clades 1 and 3 is shown on the right.

In addition, the analysis was able to identify the differences among the core and accessory genomes of *C. coli* strains from clade 1 and clade 3. *C. coli* strains from clade 1 harbored 340 core genes absent from clade 3 core genes, which in turn, harbored 159 core genes not present in clade 1 core genes (**Fig 3**). Further analysis with a higher number of *C. coli* sequences will be required in order to better estimate the core and accessory genomes of this specie. The estimated core genome for *C. jejuni* in our study (1300 genes) was close to the 1343 core genes identified in the development of the cgMLST scheme of *C. jejuni* [19] and in agreement with recent reports [33] (**S2 Table**). In addition, a significant variation in the accessory genome between *C. jejuni* and *C. coli* strains was identified, confirming the high genetic variability of *Campylobacter*. Accessory genes that are common to a particular cluster seem to contribute to modifications of carbohydrates and flagella as well as antibiotic resistance. There is also a proportion of genomes (around 28%) harboring T4SS and T6SS-related genes (see below).

### Genomic analysis of the resistome

To determine the resistome, we performed *in silico* analysis to identify genes associated with antimicrobial resistance (**Fig 4**) by screening each *Campylobacter* genome by means of the ABRicate pipeline [24] using the Resfinder, CARD, ARG-ANNOT and NCBI ARRGD databases. We additionally performed sequence alignments to identify point mutations in the *gyrA* and 23S rRNA genes. The *cmeABC* operon was present in all *C. jejuni* and *C. coli* strains analyzed. This operon encodes a common multidrug efflux pump characterized in different *Campylobacter* species, which mainly confers resistance to fluoroquinolones and in some cases to macrolides [34]. Additionally, the point mutation in the quinolone resistance-determining region (QRDR) of the *gyrA* gene, leading to the T86I substitution in GyrA, was identified in 53.1% of the strains. This mutation confers high levels of resistance to fluoroquinolones [35].

**Fig 4.**
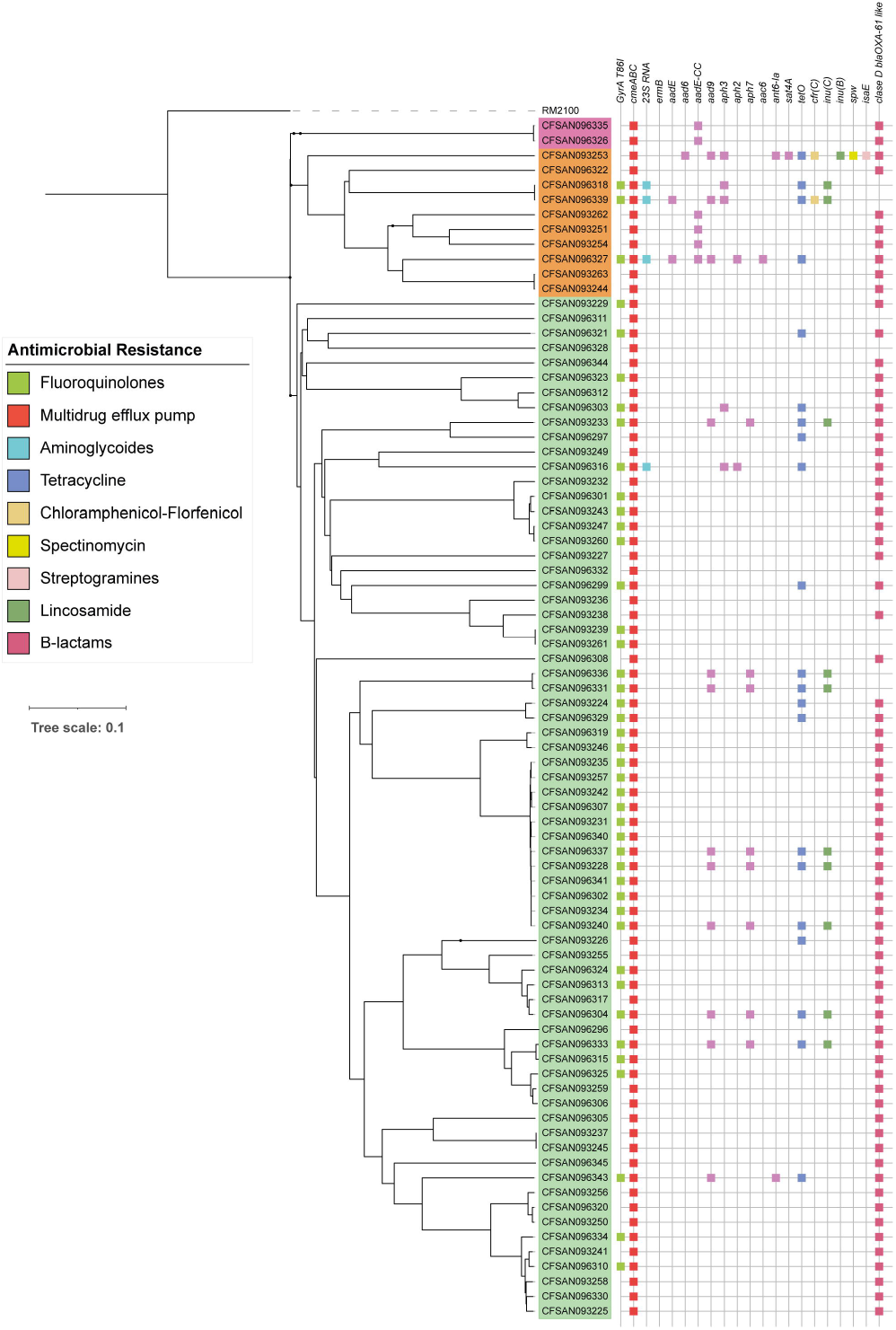
Distribution of antimicrobial resistance genes. Binary heatmaps show the presence and absence of antimicrobial resistance genes. Colored cells represent the presence of genes. cgMLST phylogenetic tree of Fig 1A is shown on the left. Strain names are color coded to highlight phylogenetic groups: *C. jejuni* strains (light green), *C. coli* clade 1 strains (orange) and *C. coli* clade 3 strains (magenta).

In contrast, a much lower fraction of the strains harbored resistance markers for other antimicrobials. *tetO* gene, which confers tetracycline resistance in *Campylobacter* spp, is found in 22.2% of the strains [36]. Furthermore, we detect the mutation of the 23S rRNA gene in 4.94% of the strains which is associated to macrolide resistance [37]. The *ermB* gene was not detected. Finally, we also found a high proportion of strains (79%) harboring the blaOXA-61 gene among *C. jejuni* and *C. coli* strains. This genetic marker is associated with β-lactam resistance and has high prevalence in *Campylobacter* strains worldwide [38].

### Presence and diversity of virulence gene content among Chilean Campylobacter strains

To determine the virulence gene content of the *C. jejuni* and *C. coli* strains, we first assembled a list of 220 potential virulence genes, including genes described in the Virulence Factor Database [39] and genes reported to contribute to the virulence of *Campylobacter* in the literature [40–43]. These genes were grouped into five distinct categories (adhesion and colonization, invasion, motility, secretion systems and toxins) and were used to screen each of the 81 genomes using BLASTn (**Fig 5** and **S1 Fig** and **S3 Table** for the full list). Most of the described factors involved in adhesion and colonization (*cadF, racR, jlpA, pdlA, dnaJ, aphC*) were prevalent just among the *C. jejuni* strains and were not associated with any particular STs (**Fig 4** and **S3 Table**).

**Fig 5.**
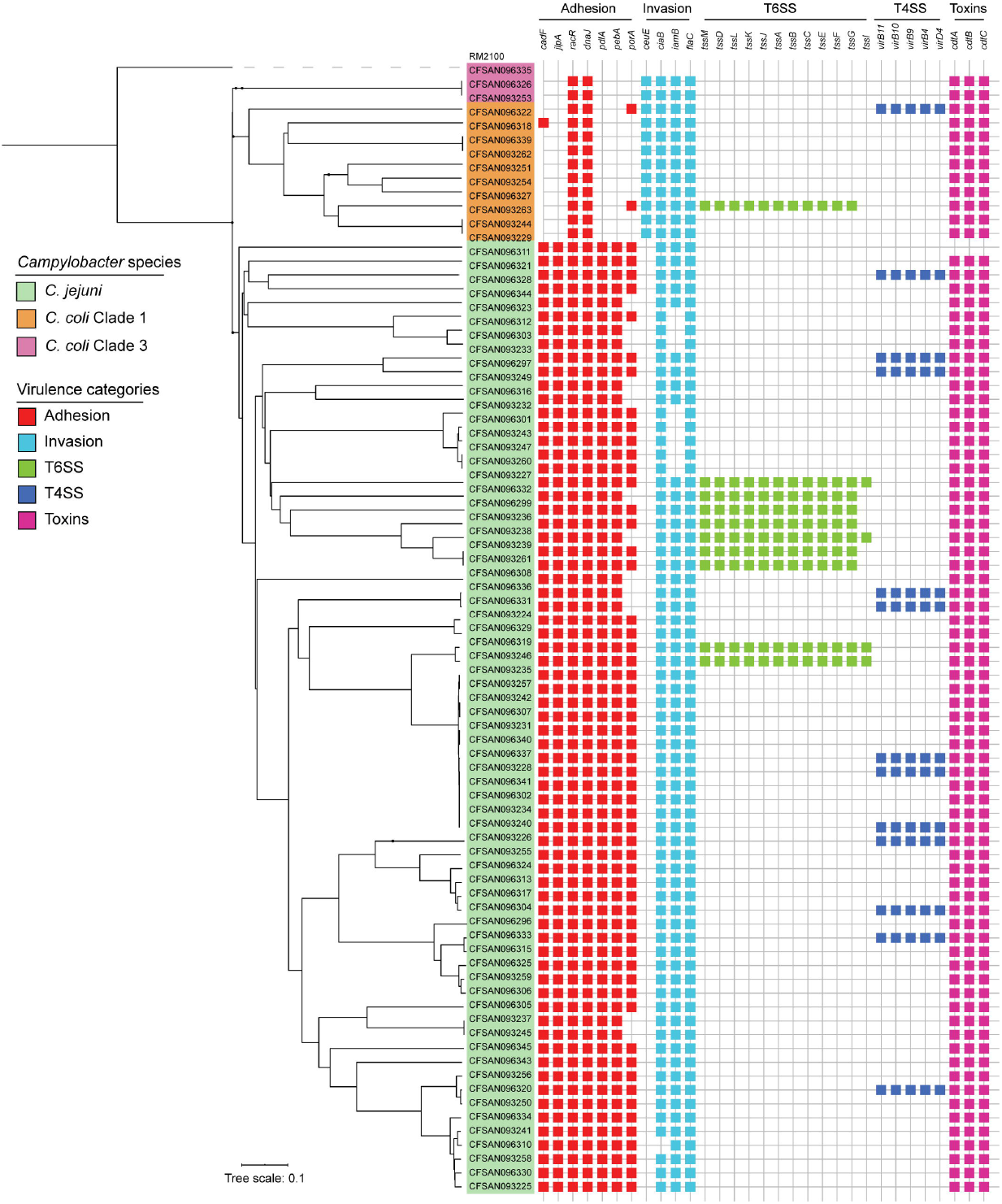
Distribution of virulence-related genes. Binary heatmaps show the presence and absence of virulence genes. Colored cells represent the presence of genes. cgMLST phylogenetic tree of Fig 1A is shown on the left. Strain names are color coded to highlight *C. jejuni* strains (green), *C. coli* clade 1 strains (orange) and *C. coli* clade 3 strains (purple).

The genes responsible for the production of the CDT toxin (*cdtA, cdtB* and *cdtC*) were also found in most strains of *C. jejuni* and in every *C. coli* strain tested. The *cdtA* and *cdtB* genes were not detected in the *C. jejuni* strain CFSAN093229. In contrast, the *ciaB* gene was identified in 98.7% of the *C. jejuni* strains (**Fig 5**). The product of this gene is involved in cecal colonization in chickens and in translocation into host cells in *C. jejuni* [44,45].

The major differences among *C. jejuni* strains were observed for genes located at the *cps* locus (Cj1413c-Cj1442) involved in capsule polysaccharide synthesis, the LOS locus (C8J1074-C8J1095) involved in lipooligosaccharide synthesis, a small set of genes in the O-linked flagellin glycosylation island (Cj1321-Cj1326) and the T6SS and T4SS gene clusters (**Fig 5** and S1 **Fig and** S3 **Table**). Additionally, *C. coli* strains carried most of the genes known to be associated with invasiveness in *C. jejuni*, including the *iamb, flaC* and *ciaB* genes. Both *C. coli* clades also harbored some genes associated with other steps of infection such as the *cdt* toxin gene and most of the flagellar biosynthesis and chemiotaxis-related genes described for *C. jejuni* (**S1 Fig**). its relevance in pathogenicity remains unclear [46].

### Distribution of the T6SS and T4SS gene clusters

Our analysis identified the T6SS gene cluster in nine *C. jejuni* strains and in one *C. coli* strain from clade 1 (**Fig 5**). As shown in **S2 Fig**, whenever present the genetic structure and the sequence identity of this cluster was highly conserved among *C. jejuni* and *C. coli* strains. For strains with genomes that were not closed, we were not able to confirm whether the *tssI* gene is located within the genetic context of the of the T6SS cluster, since it was not assembled in the same contig as the rest of the cluster. The *Campylobacter* T6SS gene cluster is often encoded within the CjIE3 conjugative element in the chromosome of *C. jejuni* [41], but it can also be found in plasmids [47,48]. Since most of our sequenced strains correspond to draft genome assemblies, we could not determine whether the T6SSs were plasmid or chromosomally encoded in most of them. However, we have closed the genomes of 2 of of these strains (CFSAN093238 and CFSAN093227) [15]. In both of them, T6SS was inserted in the chromosomal CjIE3 element, while in strain CFSAN093246, it was plasmid-encoded. Sequence-based analysis of this plasmid showed a high degree of identity with plasmid pMTVDDCj13-2 (**Fig 6**), which encodes a complete T6SS gene cluster [48].

**Fig 6.**
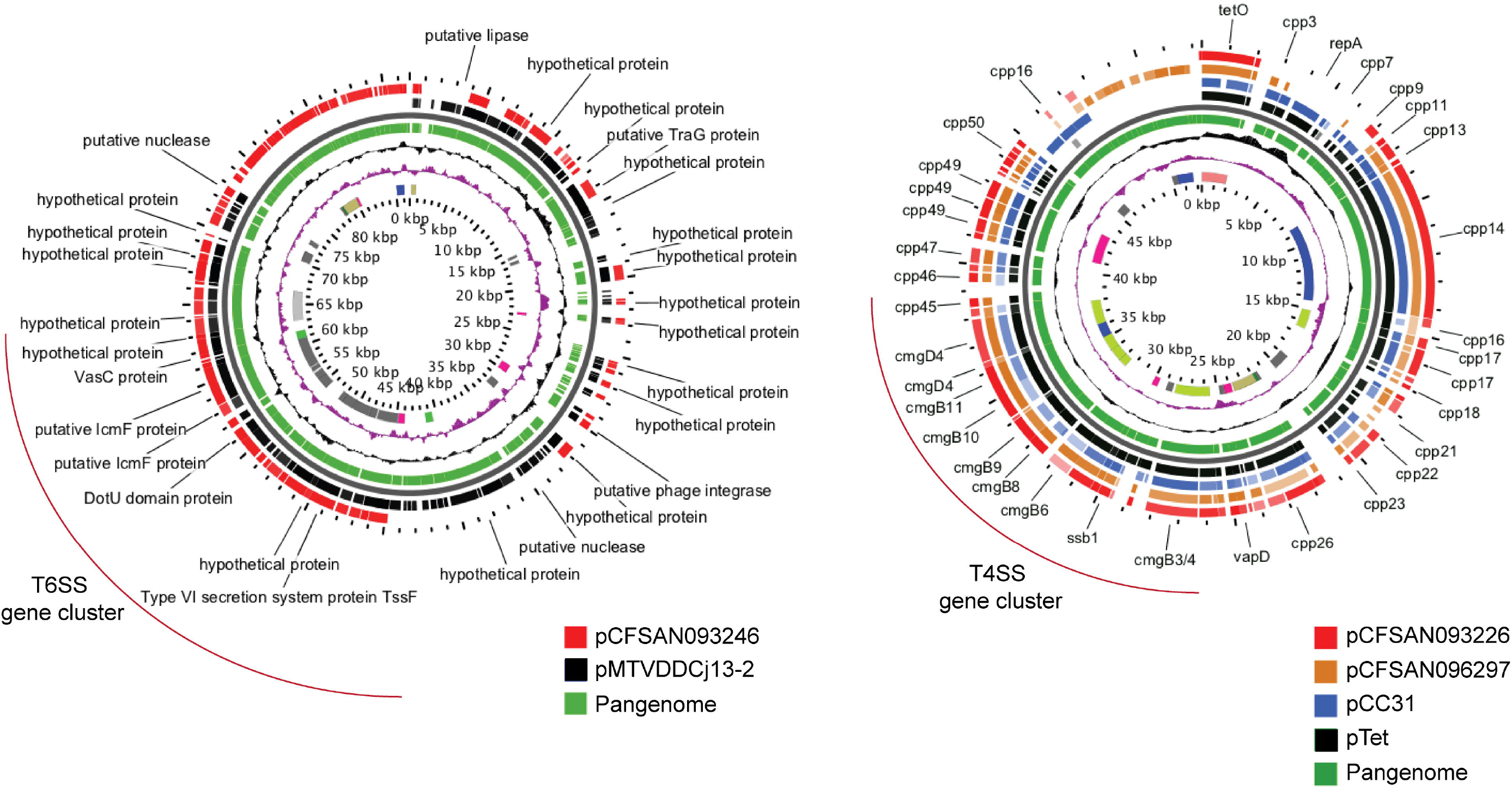
Comparative genomic analysis of plasmids encoding T6SS (A) and T4SS (B) gene clusters. Pangenome BLAST analysis was performed using the Gview server. T6SS and T4SS gene clusters are highlighted in red brackets.

Genome analysis identified a T4SS gene cluster in 12 strains of *C. jejuni* and 1 strain of *C. coli* clade 1 (**Fig 5**). It harbored the 16 core genes and was correctly assembled in individual contigs in 11 strains, which allowed the analysis of the genetic context of the cluster. In each strain, the T4SS cluster had a conserved genetic structure (**S2B Fig**). One of the main differences between the T4SS gene clusters encoded in plasmids pTet and pCC31 is the sequence divergence of the *cp33* gene [49]. While *cp33* present in strain CFSAN096321 showed high identity with the gene harbored in the pTet plasmid, every other strain possessed high identity to the *cp33* encoded in pCC31 plasmid. Both T4SS have been involved in bacterial conjugation and are not directly linked to the virulence of *C. jejuni* and *C. coli* strains [49]. As shown in **Fig 6B**, plasmids from closed genome strains CFSAN093226 and CFSAN096297 share significant similarity with both the pCC31 and pTET plasmids beyond the T4SS gene cluster. For all other strains, BLASTn analysis of each of our draft genome assemblies identified most of the plasmid genes found in pCFSAN093226, pCFSAN096297 and pTet-like plasmids, suggesting that all the T4SS gene clusters identified in these strains are encoded in plasmids as well.

## DISCUSSION

In South America, campylobacteriosis is an emerging and neglected foodborne disease. In countries such as Chile, even though *Campylobacter* is a notifiable enteric pathogen under active surveillance by public health agencies, routine stool culture–testing for this pathogen is rarely performed. This is partially explained by the high costs associated with the culture of this bacterium, including the need for special selective media and specific temperature and microaerophilic growth conditions [50]. As a consequence, there is limited genomic, clinical and epidemiological data available from the region, leaving an important knowledge gap in our understanding of the global population structure, virulence potential, and antimicrobial resistance profiles of clinical *Campylobacter* strains [1]. Here, we performed an in-depth genome analysis of the largest collection of clinical *Campylobacter* strains from Chile, which accounts for over 35% of the genome sequences available of clinical strains from South America.

cgMLST analysis provided insight into key similarities and differences in comparison to the genetic diversity reported for clinical *Campylobacter* strains worldwide. *C. jejuni* strains were highly diverse with CC-21 being the most common. CC-21 is the largest and most widely distributed CC, representing 18.9% of all *C. jejuni* strains submitted to the PubMLST database. The high prevalence of ST-1359 in our study was unexpected, since it is infrequent within the PubMLST database (0.06%, February 2020). However, ST-1359 has been recognized as a prominent ST in Israel [51]. ST-45, the second most prevalent ST worldwide (4.2%), was not identified among our strains. Previous reports from Chile identified four major CCs, including CC-21 (ST-50), CC-48 (ST-475), CC-257 (ST-257), and CC-353 (ST-353) [9,13]. Our study is consistent with these reports, except for the high prevalence of ST-1359. Interestingly, cgMLST analysis suggested that the ST-1359 strains of our study are highly similar (**Fig 1**). While this similarity might suggest that these strains could be part of an outbreak, they harbor important differences in terms of isolation date and gene content, including the presence and/or absence of different antimicrobial resistant determinants and virulence genes (including plasmids). This data suggests that these strains most likely do not represent an outbreak but correspond to highly related strains with a possible common source.

Further studies are needed to determine if ST-1359 represents an emergent ST. Interestingly, it has recently been suggested that different geographical regions within Chile harbor distinct *C. jejuni* STs. Collado *et al* described 14 STs that were exclusively identified in clinical *C. jejuni* strains isolated in the South of Chile (Valdivia) that were absent from the strains isolated from Central Chile (Santiago). Ten of these STs were also absent in our study supporting the notion that there is a differential distribution of clinically relevant STs in the country [9].

Our study also provided the first genome data of clinical *C. coli* strains from Chile. *C. coli* is divided into three genetic clades [52]. Clade 1 strains are often isolated from farm animals and human gastroenteritis cases while clade 2 and 3 strains are mainly isolated from environmental sources [52]. We showed that clade 1 strains from CC-828 were most prevalent, which is consistent with the worldwide distribution of this CC [52]. Interestingly, despite the lower amount of *C. coli* strains isolated in the 2-year period of our study (n=12), we were able to identify strains from the uncommon clade 3. Therefore, larger genomic epidemiological studies are needed to determine the prevalence of clade 3 strains in Chile and South America.

Interestingly, the genotype frequencies from clinical *Campylobacter* strains from Chile differs from the frequencies observed in other countries from South America, including Peru, Brazil and Uruguay. Clonal complex distribution that was more similar to the frequencies observed in countries such as United States and United Kingdom (Figure 2B). While it is tempting to speculate that these differences reflect distinct epidemiological scenarios, it could very well be due to the limited data currently available from South America. It has been previously noted that there is currently insufficient epidemiological data from South America to provide an accurate assessment of the burden of campylobacteriosis in the region [1]. This is reflected in the data currently available in the pubMLST database. There is information regarding only 207 clinical *Campylobacter* isolates from South America in contrast to 14,747 isolates from the United Kingdom alone. This highlights the need for larger and more representative studies. Notably, two recent genomic epidemiology studies have added valuable data from both Peru and Brazil. Pascoe *et al*., sequenced and analyzed the genomes of 62 *C. jejuni* strains of a longitudinal cohort study of a semi-rural community near Iquitos in the Peruvian Amazon [6]. They found distinct locally disseminated genotypes and evidence of poultry as an important cause of transmission. In addition, Frazao *et al*., recently sequenced and analyzed a collection of 116 *C. jejuni* strains from Brazil. The collection spanned a period of 20 years and included strains isolated from animal, human, food and environmental sources [53], identifying high levels of resistance to ciprofloxacin and tetracycline and potential transmission from nonclinical sources.

Although most *Campylobacter* infections are self-limiting, antibiotics are indicated for patients with persistent and severe gastroenteritis, extraintestinal infections, or who are immunocompromised [1]. In Chile there is little information about antimicrobial resistance levels in *Campylobacter* spp and most studies are restricted to the Central and Southern regions of the country [9–12]. The high percentage of *C. jejuni* strains with a mutation of the *gyrA* gene, which confers quinolone resistance, is consistent with recent studies from Chile, reporting ciprofloxacin resistance of 30-60% [9,11–13]. One study reported that most resistant strains were associated with ST-48 [54], but we did not find an association to any particular STs. However, all of the strains belonging to ST-1359, the most frequent ST identified, harbored this mutation. Since fluoroquinolones are not the first choice antimicrobials for human campylobacteriosis, it has been suggested that high levels of resistance might be a consequence of their broad use in animal husbandry in Latin America [55]. These animal production practices might be responsible for the dissemination of antibiotic resistance genes among Chilean *Campylobacter* strains.

Only a few clinical strains harbored the mutation of the 23S rRNA gene which confers macrolide resistance, which is consistent with their choice as first line treatment for *Campylobacter* infections [37]. Tetracycline resistance is widespread among *Campylobacter* strains worldwide [35]. Recently, the European Union reported high levels of resistance in *C. jejuni* strains (45.4%) and even higher levels in *C. coli* (68.3%). Tetracycline is especially used in the poultry industry worldwide [56], and might serve as an important reservoir for resistant strains [57]. Indeed, it was recently reported that 32.3% of *C. jejuni* strains isolated from poultry in Chile were resistant to tetracycline [12]. Our data are consistent with this report, as 22% of the clinical *C. jejuni* strains harbored the *tetO* gene. This potential emergence of resistance highlights the need for a permanent surveillance program to implement control measures for tetracycline usage in animal production.

The results of this study provide critical insights on the levels of antimicrobial resistance in *Campylobacter*, supporting previous reports that show high resistance against fluoroquinolones and tetracycline in the Chilean strains as well as high sensitivity to erythromycin [9,12]. Altogether, the data contributes to our knowledge of antimicrobial resistance which could also contribute to the development of antimicrobial surveillance programs of *Campylobacter* antimicrobial resistance in Chile.

While the ability of *Campylobacter* to cause human disease is thought to be multifactorial, there are several genes associated with the virulence of *Campylobacter*, even though their role in campylobacteriosis is not fully understood [58]. Genome analysis showed that clinical *C. jejuni* strains harbored most of the known virulence factors described for the species. On the contrary, *C. coli* strains lacked most virulence genes described for *C. jejuni* (**Fig 5**). Since most of our knowledge regarding the virulence of *Campylobacter* comes from studies performed in *C. jejuni*, this highlights the need for a better understanding of how *C. coli* strains cause disease. A limitation of our resistome and virulome analysis is that most of our data comes from draft whole genomes. Therefore, it is possible that the presence/absence of some loci may have been missed by our analysis. Nevertheless, our dataset of 17 closed genomes allowed us to gain insight into the genomic context of the T4SS and T6SS gene clusters in *C. jejuni*.

To date, there is only one T6SS cluster described in *Campylobacter* [40]. This T6SS has been shown to contribute to host cell adherence, invasion, resistance to bile salts and oxidative stress [59–61], and required for the colonization of murine [59] and avian [61] infection models. We identified a complete T6SS gene cluster in 9 *C. jejuni* and in 1 *C. coli* strain (**Fig 5**). No correlation was found between presence of the T6SS and any particular ST, which differs from what has been described in Israel, where most T6SS-positive strains belong to ST-1359 [51]. Although ST-1359 was the most common ST described in this study, none of the strains carried a T6SS.

Two distinct T4SSs have been described in *Campylobacter*. One T4SS is encoded in the pVir plasmid of *C. jejuni* and has been shown to contribute to the invasion of INT407 cells and the ability to induce diarrhea in a ferret infection model [42,62]. The second T4SS is encoded in the pTet and pCC31 plasmids of *C. jejuni* and *C. coli*, which contributes to bacterial conjugation and is not directly linked to virulence [49]. Each of the T4SS gene clusters identified in our strains showed a high degree of identity to the pTet and pCC31 T4SSs. Suggesting that they do not directly contribute to virulence, but they could facilitate horizontal gene transfer events that could lead to increase fitness and virulence.

Altogether, we provide valuable epidemiological and genomic data of the diversity, virulence and resistance profiles of a large collection of clinical *Campylobacter* strains from Chile. Further studies are needed to determine the dynamics of transmission of pathogenic *Campylobacter* to humans and the potential emergence of new virulence and antimicrobial resistance markers in order to provide actionable public health data to support the design of strengthened surveillance programs in Chile and South America.

## AUTHOR CONTRIBUTIONS

Veronica Bravo: Conceptualization, Methodology, Formal analysis, Investigation, Visualization, Data curation, Writing-Original Draft and review and editing.

Assaf Katz: Methodology, Formal analysis, Investigation, Visualization, Data Curation, Resources, Writing-Original Draft and review and editing.

Carmen Varela: Resources, Writing - Review and editing

Lorena Porte: Resources, Writing - Review and editing.

Thomas Weitzel: Resources, Writing - Review and editing.

Narjol Gonzalez-Escanola: Conceptualization, Methodology, Formal analysis, Resources, review and editing final manuscript

Carlos J Blondel: Conceptualization, Methodology, Formal analysis, Investigation, Visualization, Resources, Data curation, Writing - Review and editing, Funding acquisition, Supervision and Project administration.

## Supporting information

**S1 Fig. Distribution of the presence (red) and absence (blue) of virulence-related genes in *C. jejuni* and *C. coli* strains from this study**.

**S2 Fig. Comparative genomic analysis of T6SS and T4SS gene clusters**. (A) T6SS gene clusters of *C. jejuni* and *C. coli* strains (bold) compared to the cluster of *C. jejuni* strain 108. Genes *tagH, tssD, tssB* and *tssC* are shown in color. (B) T4SS gene clusters of *C. jejuni* strains in comparison to the T4SS gene clusters of pTet and pCC31. The *cp33* gene is highlighted in orange. BLASTn alignments were performed and visualized using EasyFig.

**S1 Table. Metadata of strains used in this study**.

**S2 Table. Frequency and distribution of CCs and STs of strains of this study**.

S3 Table. Identification of virulence genes. Cells with a red background represent presence. Empty cells represent absence of the gene in each genome.

