## Supplementary figures and images for "Genomic analysis of the diversity, antimicrobial resistance and virulence potential of clinical *Campylobacter jejuni* and *Campylobacter coli* strains from Chile"

### S1 Fig

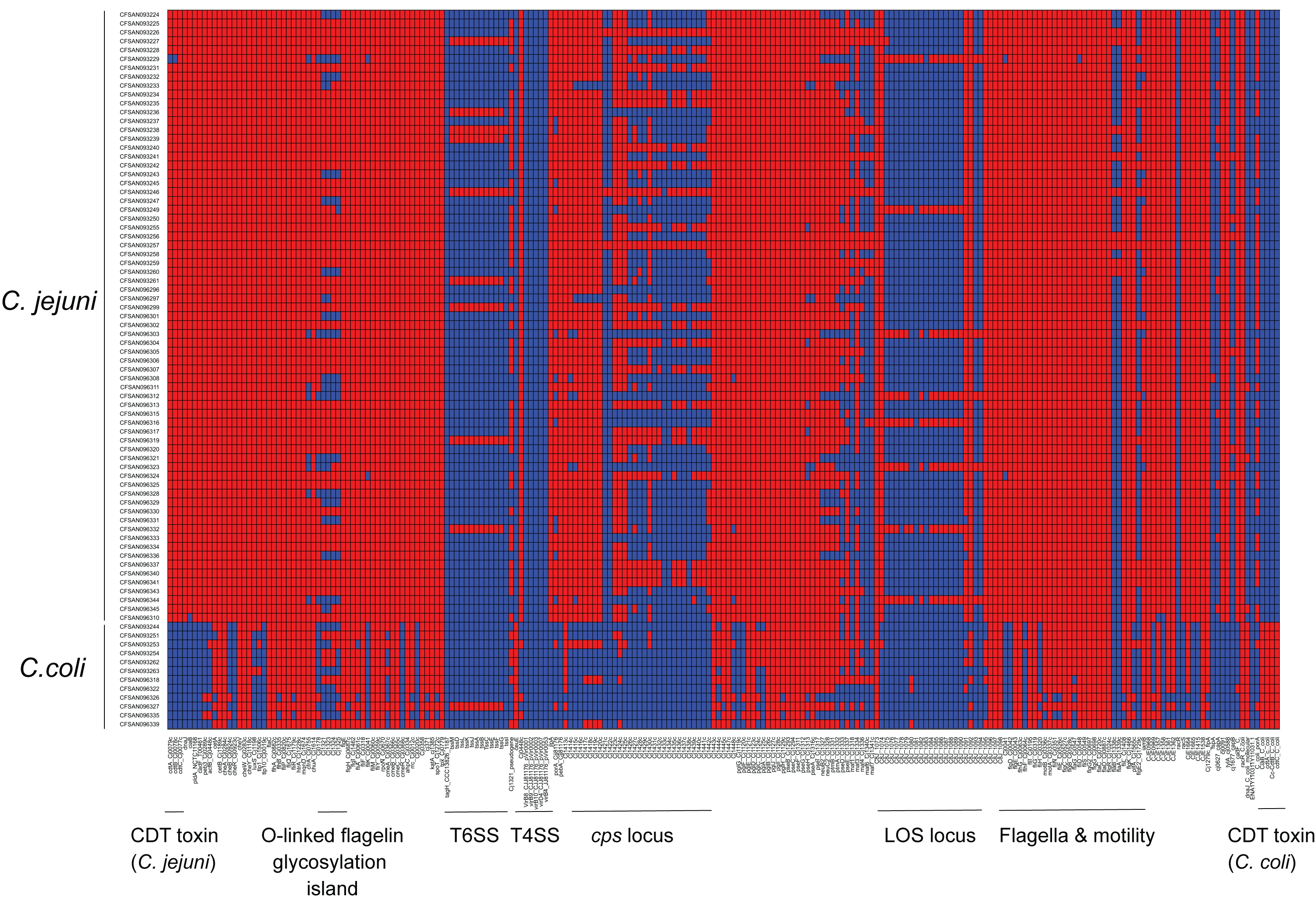

### S2 Fig

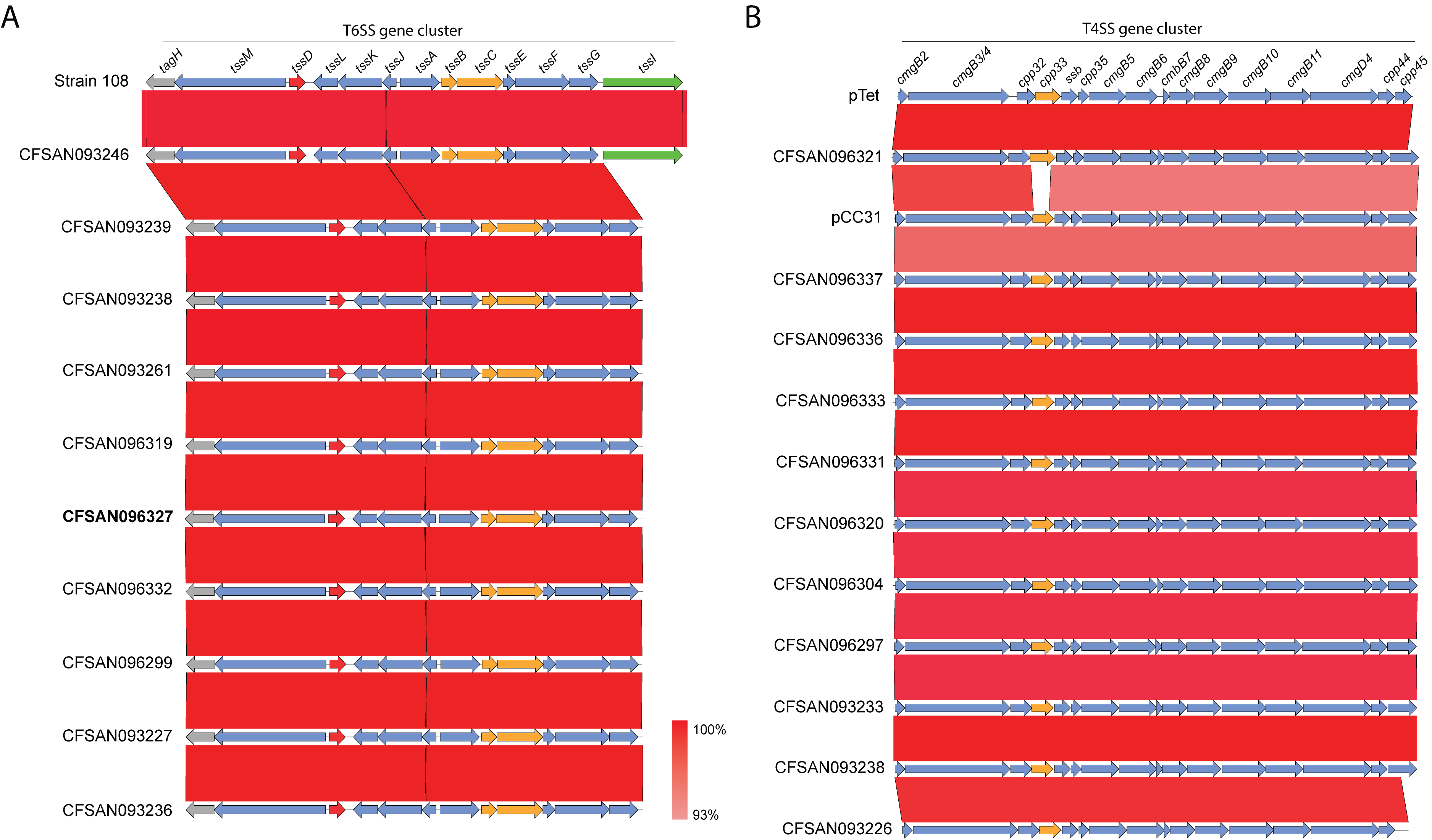
